# CellCousin2: An Optimized System for Partial Ablation and Tracing of Regenerative Lineages

**DOI:** 10.1101/2025.05.23.655316

**Authors:** Gabriel Garnik Hovhannisyan, Tawba Akhourbi, Sema Elif Eski, Isabelle Pirson, Esteban N. Gurzov, Sumeet Pal Singh

## Abstract

Regeneration can rely on multiple cellular sources, including stem cells, self-duplicating cells, and transdifferentiating cells. A central question in regenerative biology is how these distinct lineages contribute to repair and interact within a functional regenerate. We previously developed the CellCousin system to study hcellular plasticity using inducible recombination and nitroreductase-mediated ablation in zebrafish. Here, we present CellCousin2, which introduces two key improvements for long-term tracking of spared and regenerating cells. First, to reduce background recombination, we developed a Dihydrofolate Reductase (DHFR)-CreER system with dual control: DHFR-mediated degradation in the absence of trimethoprim, and tamoxifen-dependent activation. This combination minimizes leakiness while maintaining high recombination efficiency. Second, we replaced the original nitroreductase with NTR2.0, enabling effective ablation with tenfold lower metronidazole concentration, reducing off-target effects on the liver. Together, these enhancements make CellCousin2 a robust platform for dissecting the dynamics and interactions of regenerative lineages.

## Introduction

A central question in regenerative biology is how distinct cellular lineages contribute to tissue repair and how their interactions shape a functional regenerate. Regeneration can involve multiple sources, including tissue-resident stem cells, self-duplicating differentiated cells, and cells that transdifferentiate from one lineage to another. Lineage tracing has become essential to dissect the relative contributions of these cell types during regeneration and to resolve whether repaired tissue arises from differentiation of stem cells, expansion of spared cells or *de novo* fate conversion of unrelated lineages.

The cellular source of liver regeneration provides an ideal paradigm to explore the contribution of different lineages to tissue repair. In response to mild injury, liver regeneration is primarily driven by proliferation of surviving hepatocytes ^1–7^. However, in situations of extensive damage or compromised hepatocyte proliferation, cholangiocytes (biliary epithelial cells) can transdifferentiate into hepatocytes ^8–22^. This facultative regeneration mechanism has been documented in zebrafish ^19–22^, rodents ^12–18^, and even in human patients with chronic liver disease ^8–11^, highlighting the liver as a model to study lineage plasticity during regeneration.

To experimentally dissect these lineage dynamics in the liver, we previously developed the CellCousin system in zebrafish (**Fig. 1**) ^23^. This dual-reporter model is based on a Cre-inducible recombination cassette, BB-NTR, which is inspired by the Brainbow strategy ^24^. It allows stochastic expression of mutually exclusive fluorescent proteins—mTagBFP2 ^25^, histone-tagged mGreenLantern ^26^ (H2B-mGL), or mCherry ^27^—under the control of the hepatocyte-specific fabp10a promoter. In one recombination outcome, mCherry is fused to nitroreductase (NTR), resulting in expression of mCherry-NTR in a subset of hepatocytes. These mCherry-NTR– expressing cells can be selectively ablated by metronidazole (Mtz) treatment, while the H2B-mGL–expressing subpopulation is spared (**Fig. 1A**). Following mosaic ablation, the system enables tracking of spared hepatocytes and identification of *de novo* hepatocytes that arise during regeneration (**Fig. 1B**). Using CellCousin, we demonstrated that liver regeneration in late larval zebrafish involves both self-replication of spared cells and transdifferentiation from cholangiocytes ^23^.

**Figure 1:**
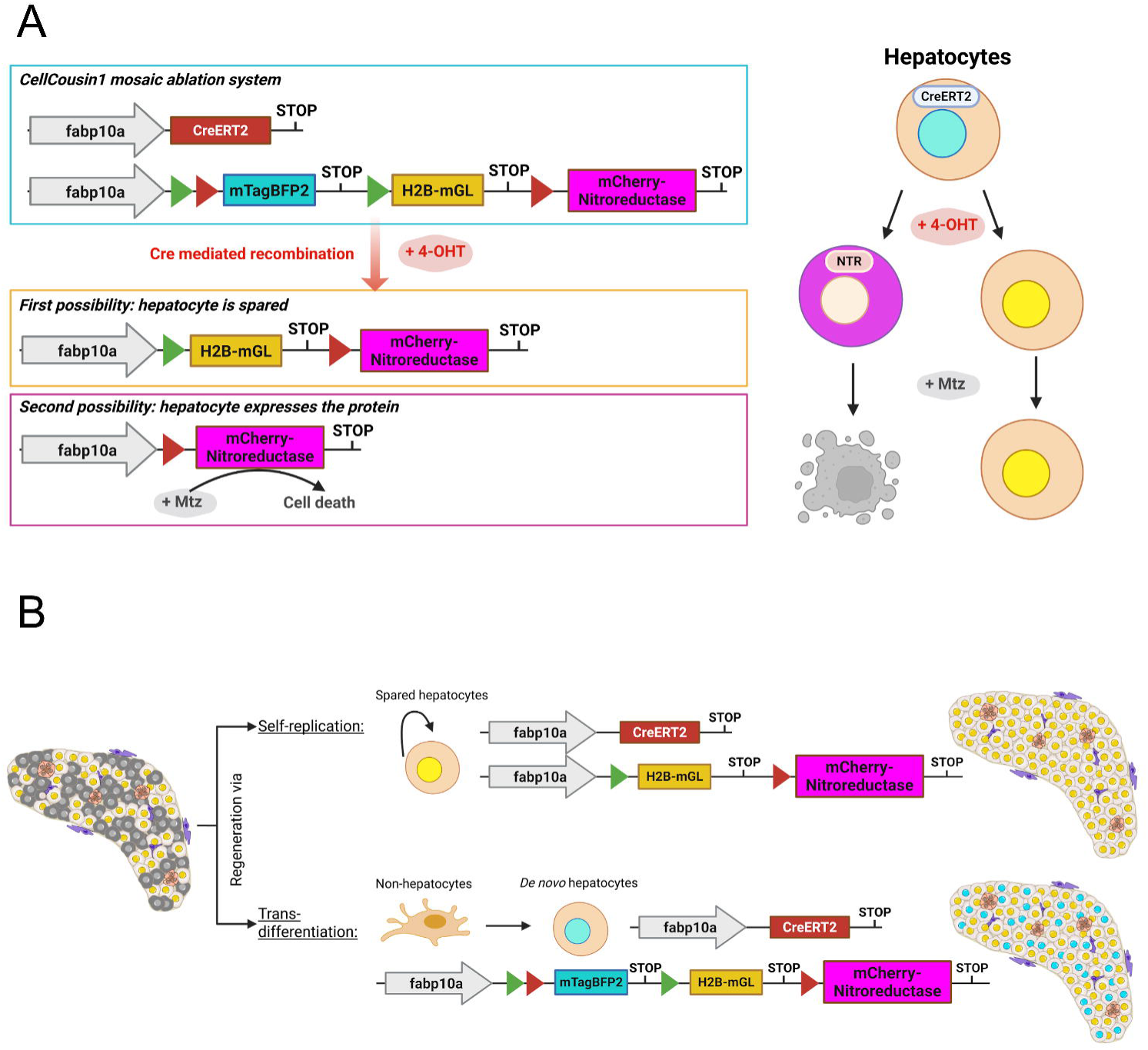
The CellCousin system for partial ablation and tracing of regenerating lineages. **(A)** Schematic overview of the CellCousin zebrafish model, which enables inducible, mosaic expression of Nitroreductase (NTR) in hepatocytes for targeted ablation. Hepatocyte-specific expression is driven by the fabp10a promoter. In the absence of recombination, hepatocytes express nuclear-localized mTagBFP2. Upon 4-hydroxytamoxifen (4-OHT) induction, Cre-mediated recombination results in the stochastic expression of either H2B-mGL (nuclear GFP) or mCherry-NTR fusion protein. Following treatment with Metronidazole (Mtz), only mCherry- NTR–expressing hepatocytes undergo cell death, allowing for controlled, partial ablation within the liver. Created in BioRender. Gurzov, E. (2025) https://BioRender.com/7dii7ih **(B)** Conceptual model of hepatocyte regeneration following ablation. Two potential regenerative routes are depicted: (1) Self-replication of spared H2B-mGL–expressing hepatocytes, and (2) Transdifferentiation of non-hepatocyte populations, giving rise to *de novo* hepatocytes that express mTagBFP2. Created in BioRender. Gurzov, E. (2025) https://BioRender.com/7qnxouj

However, the utility of the original CellCousin system is constrained by the limitations of tamoxifen-inducible CreER, particularly its background recombination in the absence of tamoxifen ^22^. In the zebrafish liver, this leakiness severely compromises temporal control, leading to premature labeling of hepatocytes. In *Tg(fabp10a:CreER)*, for instance, over 80% of hepatocytes are recombined by 2 months post-fertilization even without induction, as reported by our group and others ^22^. Such unintended recombination hinders the ability to distinguish newly generated cells from pre-labeled ones in regeneration studies.

Another limitation of the original system is the requirement for high concentrations of metronidazole (Mtz), typically 10 mM, to achieve efficient ablation of NTR-expressing hepatocytes ^28^. Prolonged exposure to such concentrations has been shown to cause off-target toxicity, potentially complicating interpretation of regenerative outcomes ^29,30^.

In this study, we present CellCousin2, an improved system for lineage tracing and mosaic ablation in the zebrafish liver. CellCousin2 incorporates two key advances: a chemically stabilized CreER recombinase that minimizes background recombination while retaining high induction efficiency, and the next-generation NTR2.0 enzyme ^31^, which enables effective ablation at tenfold lower Mtz concentrations. These enhancements make CellCousin2 a robust and precise platform for dissecting the cellular origins and dynamics of liver regeneration.

## Results and Discussion

To increase the temporal precision and reduce background activity of Cre-mediated recombination in zebrafish hepatocytes, we first generated a modified inducible recombinase construct, *Tg(fabp10a:ERCreER)*. This construct encodes a fusion of Cre recombinase with two estrogen receptor ligand-binding domains (ER), a strategy previously shown to enhance the stringency of tamoxifen-inducible systems by further restricting nuclear entry in the absence of 4-hydroxytamoxifen (4-OHT) ^32^.

We tested the recombination efficiency of *Tg(fabp10a:ERCreER)* in combination with the multicolor reporter line *Tg(fabp10a:BB-NTR)*, which labels hepatocytes with mutually exclusive fluorescent reporters following Cre activity (**Fig. 1A**). Upon 4-OHT treatment at 4 days post-fertilization (dpf), we observed successful recombination by 10 dpf, resulting in expression of either H2B-mGL or mCherry-NTR in a mosaic pattern, confirming that ERCreER is functional in the zebrafish liver (**Supplementary Fig. 1A**). However, recombination was incomplete: 25.1% ± 4.6% of hepatocytes retained expression of the default mTagBFP2 reporter (**Supplementary Fig. 1B**), indicating that a significant fraction of hepatocytes escaped recombination despite tamoxifen induction.

To address the limitations of CreER and ERCreER, we developed a CreER system incorporating a pharmacologically controlled destabilization domain (DD) that promotes proteasomal degradation of the fusion protein under basal conditions ^33,34^. The DD can be stabilized by a small-molecule, enabling chemical control through dual induction with a stabilizing compound and 4-OHT. We tested this DDCreER construct alongside CreER and ERCreER in the context of the CellCousin model to evaluate recombination efficiency and background leakiness in the zebrafish liver.

The DD is a modified version of *Escherichia coli* (*E. coli*) dihydrofolate reductase (ecDHFR), previously engineered for efficient degradation at 25°C rather than the original 37°C variant ^35^, as the higher temperature is incompatible with zebrafish physiology. This low-temperature– sensitive DD was utilized to generate the DDCreER construct, which would be degraded under physiological conditions, thereby reducing background recombination (**Fig. 2A**). In the absence of any inducers, the fusion protein is rapidly degraded and remains inactive. When larvae are treated with trimethoprim (TMP), the DD is stabilized. Under these conditions, the DDCreER protein behaves like CreER, and remains mostly cytoplasmic. Any background recombination is due to limited passive nuclear translocation of the fusion protein. Conversely, treatment with 4-OHT alone permits nuclear translocation, but only of the small fraction of fusion protein that has not yet been degraded. Notably, proteasomal degradation by the 26S proteasome is not immediate ^36^, and newly synthesized protein can accumulate briefly, leading to limited recombination. Finally, when both TMP and 4-OHT are present, the stabilized DDCreER protein is able to translocate to the nucleus and induce recombination efficiently (**Fig. 2A**).

**Figure 2:**
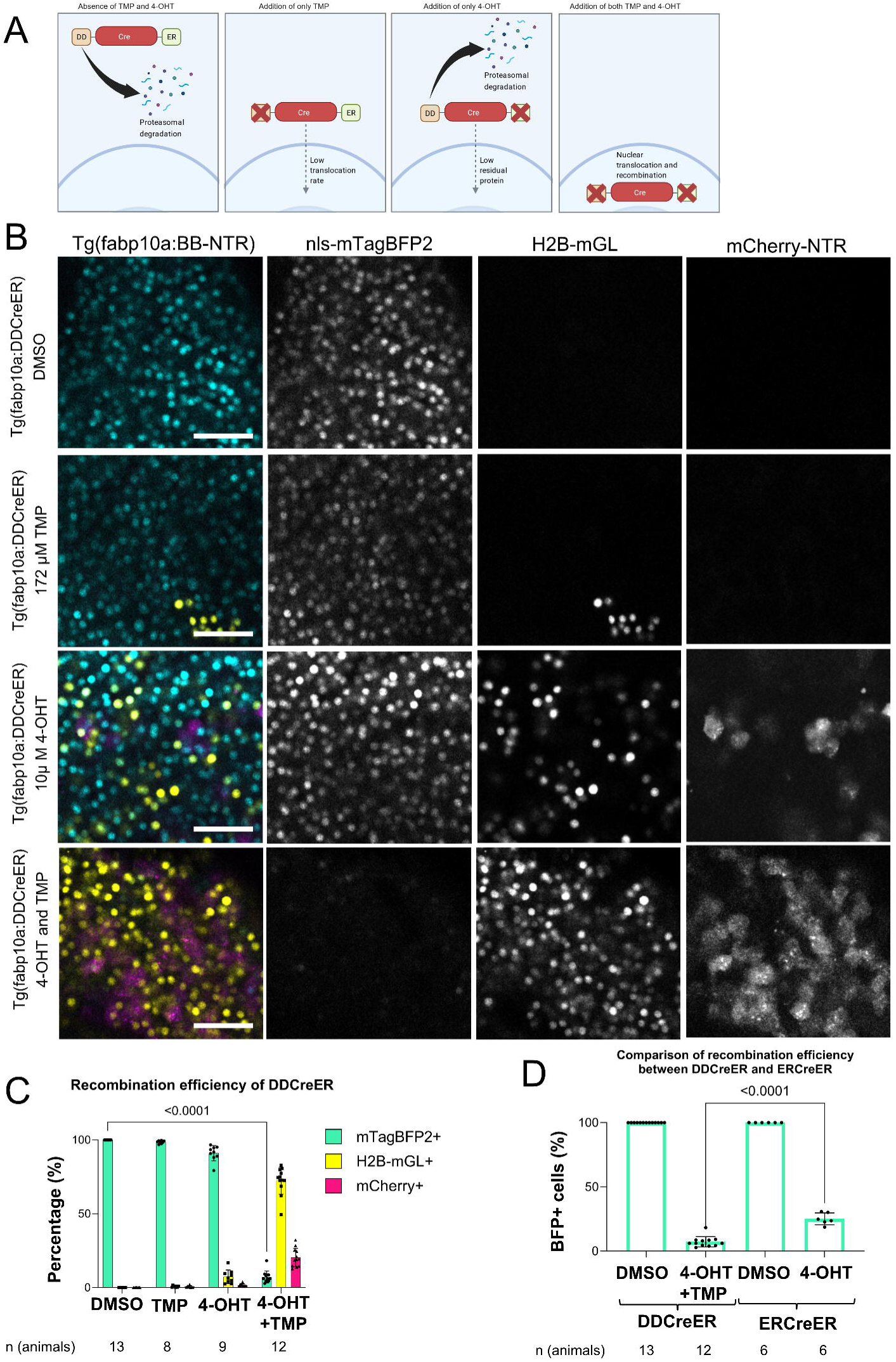
The DDCreER system enables tight control of inducible recombination in hepatocytes. **(A)** Schematic overview of the DDCreER system. In the absence of both 4-hydroxytamoxifen (4-OHT) and trimethoprim (TMP), the destabilizing domain (DD) fused CreER is targeted for proteasomal degradation. Addition of TMP alone stabilizes the DD-fused CreER, resembling the conventional CreER system and potentially allowing background recombination. Addition of 4-OHT alone promotes nuclear translocation of DDCreER, but only of the residual protein that has not yet been degraded, permitting low-level recombination. Only in the presence of both 4-OHT and TMP is DDCreER stabilized and translocated to the nucleus, enabling efficient recombination. Created in BioRender. Gurzov, E. (2025) https://BioRender.com/uteh37g **(B)** Confocal images of livers from 10 days post-fertilization (dpf) *Tg(fabp10a:BB-NTR); Tg(fabp10a:DDCreER)* zebrafish larvae following treatment with 172 µM TMP, 10 µM 4-OHT, or both together. DMSO (1%) served as a negative control. Fluorescent markers indicate non-recombined cells (mTagBFP2+), and recombined hepatocytes (H2B-mGL+ or mCherry-NTR+). Scale bar: 50 µm. **(C)** Quantification of recombination efficiency in *Tg(fabp10a:BB-NTR); Tg(fabp10a:DDCreER)* larvae under different treatments. Bar plots show the percentage of hepatocytes expressing each fluorescent marker (mTagBFP2, H2B-mGL, or mCherry). Data represents Mean ± SD. Statistical significance was calculated using two-way ANOVA followed by Tukey’s multiple comparisons test. **(D)** Comparison of recombination efficiency between the DDCreER and ERCreER systems, based on the percentage of BFP+ hepatocytes. Bar plots depict Mean ± SD. Statistical significance was assessed using ordinary one-way ANOVA followed by Tukey’s multiple comparisons test. Significantly lower proportions of unrecombined (BFP+) cells were observed in DDCreER-expressing livers.

We tested the efficiency of the new DDCreER construct by treating 4 dpf *Tg(fabp10a:DDCreER)*; *Tg(fabp10a:BB-NTR)* larvae with either 172 µM TMP, 10 µM 4-OHT or both together, and DMSO as control for 24 hours (**Fig. 2B, C**). Six days later, at 11 dpf, hepatocyte recombination was assessed by confocal microscopy. The DMSO control showed no recombination, confirming that the DDCreER construct exhibits no background activity during early development. TMP alone resulted in a recombination rate of 1.4% ± 1.0%, while 4-OHT alone yielded 9.0% ± 5.2%. The higher recombination observed with 4-OHT alone likely reflects a delay in proteasomal degradation, during which a small amount DDCreER reaches the nucleus and mediates recombination. In contrast, treatment with both 172 µM TMP and 10 µM 4-OHT led to a robust recombination efficiency of 92.8% ± 4.0% (**Fig. 2B, C**). Compared to ERCreER, which showed 25.1% ± 4.6% unrecombined nls-mTagBFP2+ cells, DDCreER achieved significantly higher recombination, with only 7.2% ± 4.0% unrecombined hepatocytes (**Fig. 2D**).

Next, to assess background recombination over long time periods, we examined livers from 2-month-old zebrafish expressing *Tg(fabp10a:BB-NTR)* in combination with either CreER, ERCreER, or DDCreER (**Fig. 3A, B**). Importantly, no 4-OHT treatment was administered in these conditions, allowing us to quantify baseline recombination leakiness. As expected, the *Tg(fabp10a:CreER)* line showed extensive background recombination, with 80.6% ± 5.4% of hepatocytes labeled. In contrast, *Tg(fabp10a:DDCreER)* showed markedly reduced background recombination, with only 6.1% ± 7.0% of hepatocytes recombined. Notably, *Tg(fabp10a:ERCreER)* exhibited negligible background activity, with a recombination rate of 0.0% ± 0.01% (**Fig. 3B**).

**Figure 3:**
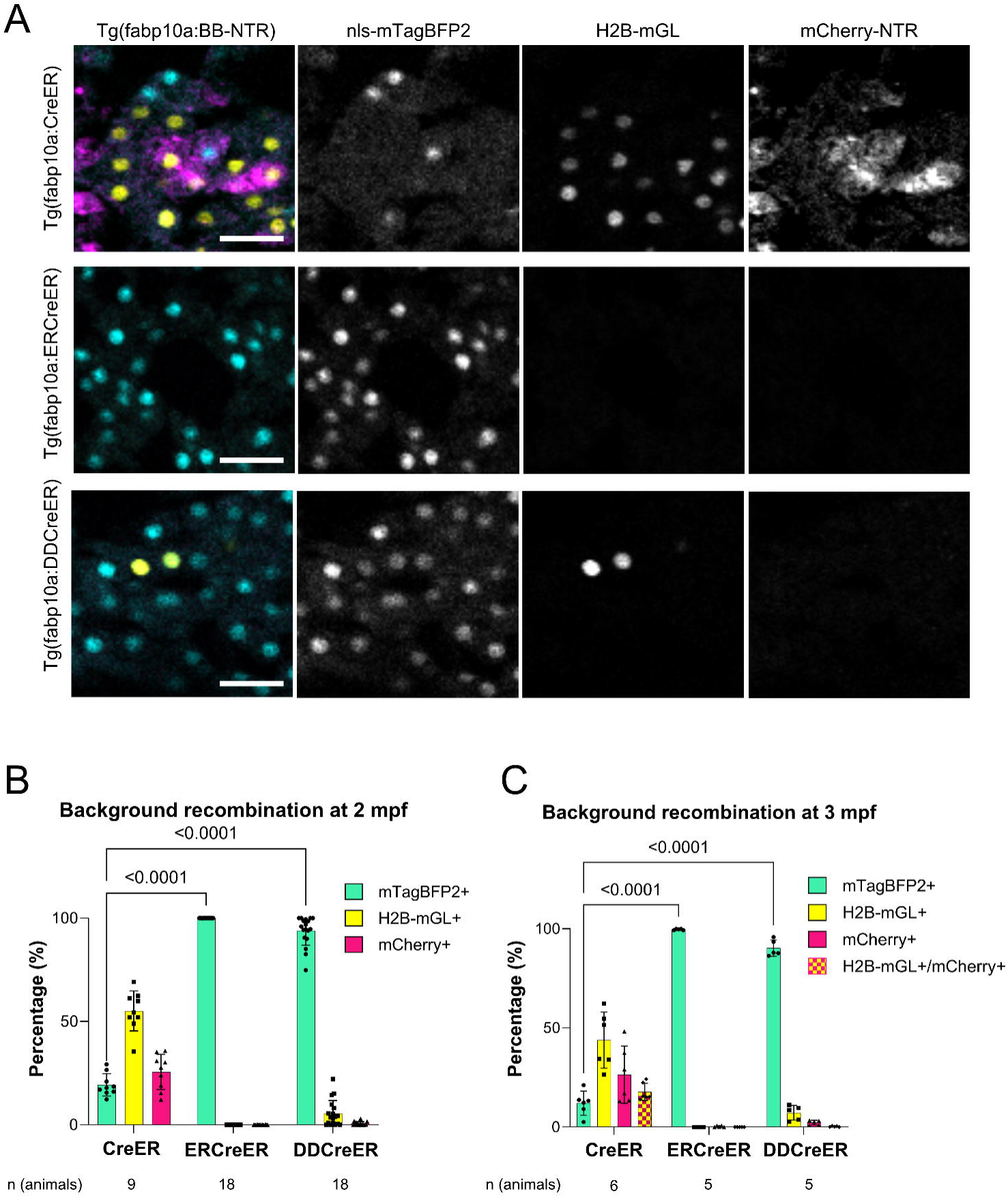
Background recombination in the CellCousin model. **(A)** Confocal images of liver cryosections from 2-month post-fertilization (mpf) *Tg(fabp10a:BB-NTR)* zebrafish expressing *CreER*, *ERCreER*, or *DDCreER*, without any 4-hydroxytamoxifen (4-OHT) or trimethoprim (TMP) treatment. Fluorescent markers indicate non-recombined nls-mTagBFP2+, or recombined H2B-mGL+ or mCherry-NTR+ hepatocytes. Scale bar: 20 µm. **(B)** Bar plot representing Mean ± SD showing quantification of background recombination at 2 mpf based on histological analysis of liver sections from *Tg(fabp10a:NTR)* zebrafish. Significance determined by two-way ANOVA followed by Tukey’s multiple comparisons test. **(C)** Flow cytometry-based quantification of background recombination at 3 mpf in *Tg(fabp10a:BB-NTR)* zebrafish. The proportion of mTagBFP2+, H2B-mGL+, mCherry+, and double-positive (H2B-mGL+/mCherry+) hepatocytes was measured by FACS. Each dot represents one animal and the bar plots represent Mean ± SD. Statistical analysis performed using two-way ANOVA followed by Tukey’s multiple comparisons test.

To independently validate these findings, we performed fluorescence-activated cell sorting (FACS) on livers from the same three genotypes at 3 months post-fertilization (**Supplementary Fig. 2A, B**). Consistent with the histological analysis, *Tg(fabp10a:CreER)* showed a background recombination rate of 88.0% ± 6.1%, whereas *Tg(fabp10a:DDCreER)* showed a substantially lower rate of 9.7% ± 4.1%. Again, *Tg(fabp10a:ERCreER)* showed minimal leakiness, with a background recombination rate of 0.3% ± 0.3% (**Fig. 3C**).

Having established the high recombination efficiency and low background activity of the DDCreER system, we next attempted to improve the efficiency of hepatocyte ablation.

The CellCousin model enables partial ablation by utilizing the codon optimized nitroreductase (NTR) derived from *E. coli* ^37–39^. Treatment with 10 mM metronidazole (Mtz) leads to the production of a DNA cross-linking agent in the NTR expressing cells, which induces cell death^28^. However, this high concentration of Mtz is associated with non-negligible off-target effects ^29,30^, potentially confounding interpretation of regenerative responses. Reducing Mtz exposure would therefore improve the physiological relevance of the system by minimizing unintended toxicity. Recently, an improved nitroreductase, NTR2.0, was isolated from the bacterial species *Vibrio vulnificus* ^31^. NTR2.0 enables efficient ablation at significantly lower Mtz concentrations. To implement this improvement, we replaced the original NTR with NTR2.0 in the BB-NTR cassette, generating the BB-NTR2 construct and transgenic line for hepatocytes.

We tested the ablation efficiency of the new *Tg(fabp10a:BB-NTR2)* line by treating larvae with different concentrations of Mtz (**Fig. 4**). To induce recombination, *Tg(fabp10a:BB-NTR2)*; *Tg(fabp10a:DDCreER)* larvae were treated at 4 dpf with 172 µM TMP and 10 µM 4-OHT. This resulted in high recombination efficiency, with only 3.8% ± 1.3% unrecombined hepatocytes, compared to 99.9% ± 0.01% in DMSO-treated controls (**Fig. 4A, B, C**).

**Figure 4:**
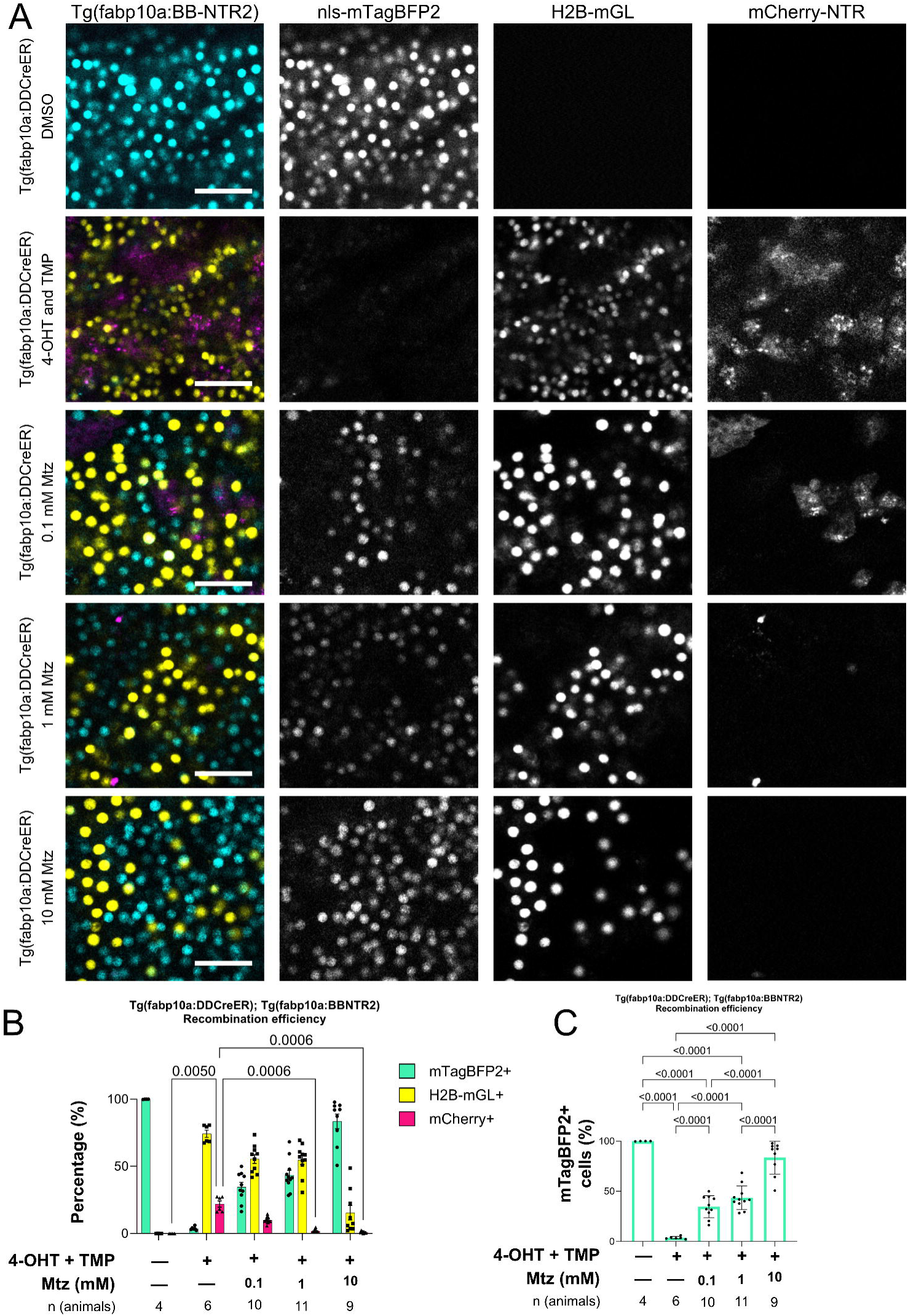
CellCousin2 enables partial hepatocyte ablation and segregation of spared and *de novo* hepatocytes. **(A)** Confocal images of livers from *Tg(fabp10a:BB-NTR2); Tg(fabp10a:DDCreER)* zebrafish larvae treated with DMSO (control) or recombination-induced using 172 µM trimethoprim (TMP) and 10 µM 4-hydroxytamoxifen (4-OHT) at 4–5 days post-fertilization (dpf). Hepatocyte ablation was performed at 10 dpf using 0.1, 1, or 10 mM metronidazole (Mtz), and livers were imaged at 14 dpf. Scale bars: 50 µm. **(B)** Bar plots showing quantification of recombination and ablation efficiency in *Tg(fabp10a:BB-NTR2); Tg(fabp10a:DDCreER)* livers. The proportion of hepatocytes expressing mTagBFP2, H2B-mGL, or mCherry-NTR2 was quantified before ablation (10 dpf) and after ablation (14 dpf) across Mtz concentrations. Data represents Mean ± SD. Significance was assessed using two-way ANOVA followed by Šídák’s multiple comparisons test. **(C)** Quantification of mTagBFP2+ hepatocytes in the liver as a proxy for unrecombined cells before ablation and newly generated hepatocytes (*de novo*) after ablation. A reduction in mTagBFP2+ cells post-recombination followed by a partial reappearance after Mtz treatment suggests regeneration via transdifferentiation. Bar plots represent Mean ± SD. Statistical significance was calculated using one-way ANOVA followed by Tukey’s multiple comparisons test.

Following recombination, larvae were treated with 0.1, 1, or 10 mM Mtz for 24 hours at 10 dpf to assess ablation efficiency. At 14 dpf, the 0.1 mM Mtz treatment caused only a modest reduction in the proportion of mCherry-NTR2+ hepatocytes (9.9% ± 2.5%), compared to pre-ablation levels (21.9% ± 5.6% at 10 dpf) (**Fig. 4B**). In contrast, both 1 mM and 10 mM Mtz treatments led to efficient ablation, reducing the mCherry-NTR2+ population to 1.8% ± 1.2% and 1.0% ± 0.7%, respectively (**Fig. 4B**). In both the 0.1 and 1 mM Mtz conditions, we observed a concomitant increase in mTagBFP2+ hepatocytes to 34.6% ± 10.9% and 43.5% ± 11.7%, respectively, suggesting regeneration by *de novo* cells (**Fig. 4A, C**). These results indicate that 1 mM Mtz is sufficient for effective ablation in the BB-NTR2 line, representing a tenfold reduction in required Mtz concentration relative to the original system.

We also noted that ablation with 10 mM Mtz appeared to induce widespread hepatocyte death, potentially due to a bystander effect, in which cytotoxic intermediates generated in dying NTR- expressing cells diffuse into neighboring cells and cause collateral damage ^40,41^. In this context, *de novo* mTagBFP2+ hepatocytes comprised 83.5% ± 16.3% of the liver, accompanied by a marked reduction in H2B-mGL+ hepatocytes, which represent the non-NTR-expressing population (**Fig. 4B, C**).

We refer to the optimized combination of *Tg(fabp10a:BB-NTR2)* and *Tg(fabp10a:DDCreER)* as CellCousin2.

We also evaluated whether the ERCreER system could be used with the newly generated *Tg(fabp10a:BB-NTR2)* line to achieve near-complete recombination efficiency and serve as a viable alternative. In initial experiments *Tg(fabp10a:ERCreER)*; *Tg(fabp10a:BB-NTR)* larvae treated with 10 µM 4-OHT for 24 hours at 4 dpf showed a sub-optimal recombination rate of 74.9% ± 4.6% (**Supplementary Fig. 1B**). To increase this rate, we tested either higher concentrations or repeated dosing of 4-OHT. A single treatment with 20 µM 4-OHT for 24 hours at 4dpf led to a recombination rate of 59.3% ± 3.9%, while a single treatment with 20 µM 4-OHT for 8 hours at 5 dpf led to a survival rate of 40% and a recombination rate of 36.4% ± 8.2% (**Supplementary Fig. 3A, B**). A double treatment, by giving one 10 µM 4-OHT treatment on 4 dpf and a second treatment on 5 dpf, reduced larval survival to 30% and resulted in a recombination rate of only 42.3% ± 7.4% (**Supplementary Fig. 3A, B**). A similar double dose of 20 µM 4-OHT resulted in 100% mortality.

Overall, increasing the 4-OHT concentration or dosing frequency seemed to paradoxically decrease recombination efficiency. Higher 4-OHT exposure led to a decrease in hepatocyte number (**Supplementary Fig. 3C**), suggesting that tamoxifen treatment may have toxic effects on the liver. Hepatocyte loss might trigger a regenerative response from non-hepatocytes via transdifferentiation, thereby increasing the proportion of mTagBFP2+ cells.

Thus, although the *Tg(fabp10a:ERCreER)* line exhibits no detectable background recombination (**Fig. 3B, C**), its increased stringency results in sub-optimal recombination in hepatocytes.

In conclusion, *Tg(fabp10a:DDCreER); Tg(fabp10a:BB-NTR2)*, termed as CellCousin2, provides an optimal balance between recombination efficiency and temporal precision, combining low background activity with robust inducibility to enable controlled mosaic labeling and ablation in the zebrafish liver. Our development of CellCousin2 provides a powerful tool for uncovering the cellular logic of liver regeneration. By enabling temporally controlled, low-background lineage labeling and selective ablation using low-toxicity conditions, CellCousin2 makes it possible to track both spared and de novo hepatocytes with high fidelity. These improvements open new opportunities for studying how cell fate decisions are coordinated during organ repair and for investigating regenerative plasticity in other tissues. Beyond the liver, the modular design of CellCousin2 can be readily adapted to other organs or transgenic models, making it a valuable resource for the regeneration and developmental biology communities.

## Methods

### Zebrafish lines and husbandry

Wild-type and transgenic zebrafish from the outbred AB strain were utilized in all experiments. Zebrafish larvae were kept at 28.5 °C, with ∼50 larvae per 30 ml of E3 medium (5 mM NaCl, 0.17 mM KCl, 0.33 mM CaCl2, 0.33 mM MgSO4, 10 mM HEPES) until 5 days post-fertilization (dpf). At 5 dpf, the larvae were fed daily with rotifers (100 µl/larva/day) and maintained at a density of around 30 larvae per 500 ml of 5 ppt fish water until 10 dpf. Subsequently, they were transferred to tanks with water flow and fed a dry fish flake diet twice daily. All zebrafish husbandry and experimental procedures for transgenic lines were conducted in accordance with institutional (Université Libre de Bruxelles) and national ethical and animal welfare guidelines and regulations, which were approved by the Ethical Committee for Animal Welfare (CEBEA) from the Université Libre de Bruxelles (protocols 864N, 865N, 877N, 881N, 882N).

In this study, the following published transgenic lines were used: *Tg(fabp10a:creERT2; cryaa:CFP)^ulb34^* and *Tg(fabp10a:lox2272-loxp-nls-mTagBFP2-stop-lox2272-H2B-mGL-stop-loxp-mCherry-NTR; cryaa:mCherry)^ulb33^* abbreviated as *Tg(fabp10a:BB-NTR)* ^23^.

### Generation of the Tg(fabp10a:ERCreER)^ulb42^, Tg(fabp10a:DDCreER)^ulb43^ and Tg(fabp10a:BB-NTR2)^ulb41^ lines

The fabp10a:ER^T2^CreER^T2^; cryaa:mCerulean construct [abbreviated as fabp10a:ERCreER] was generated by replacing the CreER^T2^ in fabp10a:CreER^T2^; cryaa:mCerulean construct (Addgene plasmid # 230044 ; http://n2t.net/addgene:230044 ; RRID:Addgene_230044) ^23^ with ER^T2^CreER^T2^ using EcoRI/NotI. The ER^T2^CreER^T2^ sequence was obtained fom pCAG-ER^T2^CreER^T2^, which was a gift from Connie Cepko (Addgene plasmid # 13777 ; http://n2t.net/addgene:13777 ; RRID:Addgene_13777) ^32^.

For the fabp10a:DDCreER^T2^; cryaa:mCerulean construct [abbreviated as fabp10a:DDCreER], the destabilisation domain (DD) sequence was based on a low-temperature–sensitive Escherichia Coli dihydrofolate reductase (ecDHFR) mutant previously described by Cho et al. ^35^. This mutant includes multiple point mutations (N23S, V78A, E120G, E134G, E154V, E157G) and enables efficient proteasomal degradation at 25°C. The low-temperature DHFR (ltDHFR) sequence was obtained from pBMN ltDHFR(DD)-YFP plasmid, a gift from Thomas Wandless (Addgene plasmid # 47076 ; http://n2t.net/addgene:47076 ; RRID:Addgene_47076). The sequence was codon optimized for zebrafish using iCodon ^42^ and fused to the CreER^T2^ sequence. The dsDNA fragment containing *ltDHFR(DD)-CreERT2*, flanked by EcoRI and NotI sites, was synthesized by GenScript Biotech and used to replace CreER^T2^ in the fabp10a:CreER^T2^; cryaa:mCerulean construct ^61^ using restriction enzyme based cloning with EcoRI/PacI following by ligation with T4 ligase.

To generate the fabp10a:lox2272-loxp-nls-mTagBFP2-stop-lox2272-H2B-mGL-stop-loxp-mCherry-NTR2; cryaa:mCherry construct [abbreviated as fabp10a:BB-NTR2], the dsDNA sequence of BB-NTR2 flanked with EcoRI/PacI was synthesized by GenScript Biotech and used to replace BB-NTR in the fabp10a:BB-NTR; cryaa:mCherry construct (Addgene plasmid #230043 ; http://n2t.net/addgene:230043 ; RRID:Addgene_230043) ^23^ using restriction enzyme based cloning with EcoRI/PacI followed by ligation with T4 ligase.

To generate transgenic lines, a solution containing the 20 ng/µl of the construct was mixed with I-SceI meganuclease enzyme and injected into one-cell stage embryos to facilitate transgenesis. Transgenic founders were identified and maintained using the eye marker expression.

### Trimethoprim (TMP) and 4-Hydroxytamoxifen (4-OHT) labeling

For consistent recombination efficiency across the experiments, appropriate concentration of 4-OHT (MedChemExpress, HY-16950) was prepared from 10 mM stock in 100% DMSO and heated at 65 °C for 10 minutes to activate trans-4-OHT ^43^. TMP (Sigma-Aldrich, T7883) stock at 172 µM was prepared by diluting 10 mg of TMP powder and 2 mL 100% DMSO in 200 mL of E3 medium, and put under agitation for 1 hour. All treatments were carried out in the dark at 28.5 °C in 30 ml E3 medium with 30 larvae per petri dish. The labeling experiments were performed under the following treatment conditions:

*Tg(fabp10a:DDCreER); Tg(fabp10a:BB-NTR)* or *Tg(fabp10a:DDCreER); Tg(fabp10a:BB-NTR2):* First treatment of 172 µM TMP at 4 dpf for 2 hours, followed by 172 µM TMP and 10 µM 4-OHT treatment for 24 hours.

*Tg(fabp10a:ERCreER); Tg(fabp10a:BB-NTR)*: 10 µM 4-OHT treatment at 4 dpf for 24 hours.

*Tg(fabp10a:ERCreER); Tg(fabp10a:BB-NTR2)*: 20 µM 4-OHT treatment at 4 dpf for 24 hours, or 20 µM 4-OHT treatment at 5 dpf for 8 hours, or 10 µM 4-OHT treatment in two separate 12-hour intervals. The first treatment was started at 4 dpf and the second treatment was performed at 5 dpf. In between the two treatments, animals were kept in E3 medium.

### Genetic partial ablation of hepatocytes

For partial ablation experiments, 10 dpf zebrafish larvae were treated with freshly prepared 0.1, 1, or 10 mM metronidazole (Mtz) (Thermo Scientific, #443-48-1) solution in 1% DMSO in E3 medium. The solution was vortexed continuously for 10 minutes to ensure complete dissolution of Mtz. Larvae were exposed for 16-hours by transferring 30 larvae to a 90 mm petri dish containing 30 ml of the Mtz solution, protected from light. A 1% DMSO solution was used as the vehicle for all control animals. After ablation, larvae were rinsed three times with E3 medium and then transferred to tanks with water flow and maintained on a dry food regime.

### Imaging

Before *in vivo* confocal imaging, larvae were anesthetized with 0.02% tricaine (MS-222) (Sigma-Aldrich, E10521) for 1 minute and then mounted in 1% Low-Melt Agarose (Lonza, 50080) containing 0.02% tricaine. Imaging was performed on a glass-bottomed dish FluoroDish™ (WPI, FD3510-100) using Zeiss LSM 780 confocal microscope platform. Livers were imaged using a 20x/0.8 air or 25x/0.8 water immersion correction lens. The imaging frame was set to 1024 × 1024 pixels. Samples were excited at 405 nm for mTagBFP2, 488 nm for mGreenLantern, and 543 nm for mCherry with fluorescence collected in the respective ranges of 426-479 nm, 497-532 nm, 550-633 nm, and 641-735 nm.

For 2 mpf imaging, zebrafish were euthanized in 0.2% Tricaine (MS-222, Sigma E10521) solution. The animals were fixed by immersion in 4% paraformaldehyde (PFA) (Thermo, J61899.AK) + 1% Triton X (Sigma, T8787) overnight at 4°C. The animals were washed 2–3 times in PBS to remove PFA before proceeding. The liver were dissected using Dumont #5 forceps (Fine Science Tools 11295-10). The tissue was immersed in 30% sucrose solution overnight at 4°C, embedded in Tissue Freezing Medium (Leica 14020108926) and frozen at −80°C. Thin sections (8 µm) were obtained using cryostat (Leica CM3050 S), collected on frosted glass slides (Thermo Scientific 12362098), and covered with glass coverslip of #1 thickness (Carl Roth GmbH NK79.1) using Fluoromount-G™ mounting medium (Invitrogen, 00-4958-02). Images were taken on Zeiss LSM 780 confocal microscope using 20x/0.8 air lens.

### Image analysis

Image analysis was performed using Imaris 10.2 (Oxford Instruments). The Imaris software was utilized to calculate liver volume by manually selecting liver segments using the ‘surface’ module.

Quantifications related to the number of hepatocytes in the *CellCousin* model were performed by manually thresholding the relevant fluorescent signal (nuclear or cytoplasmic) and separating objects based on intensity. The ‘spots’ module was used with an estimated diameter set to 6 µm for mTagBPF+ and H2B-mGL+ cells (nuclear signal), or 8 µm for mCherry+ (cytoplasmic signal).

### Statistical analysis

Statistical analysis was performed using GraphPad Prism software (version 10.2 GraphPad Software, San Diego, CA). When comparing multiple groups, we used a two-way ANOVA test followed by a multiple comparison test. No data was excluded from analysis. Blinding was not performed during analysis.

### Data Availability

Plasmids generated in this manuscript have been deposited to Addgene: fabp10a:lox2272-loxp-nls-mTagBFP2-stop-lox2272-H2B-mGL-stop-loxp-mCherry-NTR2.0; cryaa:mCherry: Plasmid #230075; fabp10a:ERT2CreERT2; cryaa:mCerulean: Plasmid #230077; fabp10a:ERT2CreERT2; cryaa:mCerulean: Plasmid #230076. Raw images and image analysis files are available upon request to the corresponding author.

## Acknowledgements

We thank the members of IRIBHM Fish Facility, Christine Dubois from FACS facility, and M Martens and JM Vanderwinden from the Light Microscopy Facility for technical assistance at ULB. ENG is a Research Associate of the Fonds de la Recherche Scientifique (FNRS), Belgium. The work was supported by FNRS grants 40021615 (FRIA) to GGH, 40006730 (ASP) to SEE,, and 40005588 (MISU-PROL), 40013427 (CDR), 40027730 (CDR) and 40020360 (PDR) to SPS, and funding from Université libre de Bruxelles, Jaumotte-Demoulin Foundation, and Ramalingaswami Re-entry Fellowship from Department of Biotechnology (DBT), India to SPS.

## Author Contributions

GGH: investigation, visualization, methodology, writing—original draft, review, and editing. TA, SEE: investigation, methodology and resources.

IP: resources.

ENG: supervision, methodology, and writing—review and editing.

SPS: conceptualization, supervision, funding acquisition, project administration, and writing— original draft, review, and editing.

