## Supplementary Figures 1-3 for "CellCousin2: An Optimized System for Partial Ablation and Tracing of Regenerative Lineages"

### Table of Contents

Supplementary Figure 1: Recombination efficiency using *Tg(fabp10a:ERCreER)*. 3

Supplementary Figure 2: Flow cytometry analysis of background recombination in hepatocytes from different transgenic zebrafish lines. 4

Supplementary Figure 3: High concentrations or prolonged exposure to 4-OHT do not enhance ERCreER recombination efficiency. 5

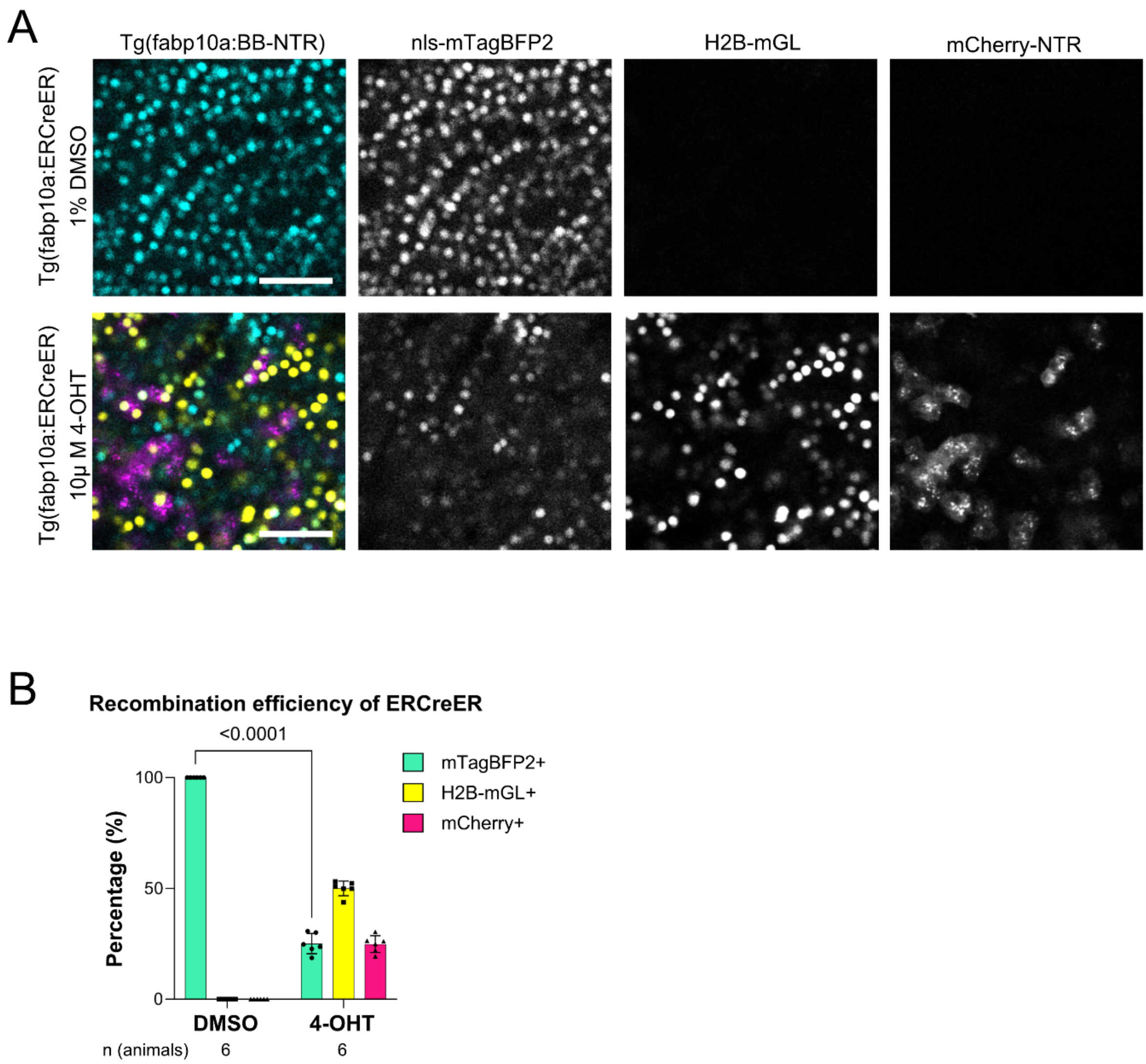


**Supplementary Figure 1: Recombination efficiency using *Tg(fabp10a:ERCreER)*.**

**(A)** Confocal images of livers from 10 dpf *Tg(fabp10a:BB-NTR)*; *Tg(fabp10a:ERCreER)* zebrafish larvae. Animals were treated with 1 % DMSO (control) or 10 µM 4-OHT from 4 – 5 dpf. Scale bar: 50 µm. **(B)** Barplot with Mean±SD showing percentage of hepatocytes expressing specific fluorescent proteins in DMSO or 4-OHT treated groups (n=6 animals). Significance calculated with Two-way ANOVA followed by Šídák's multiple comparisons test.

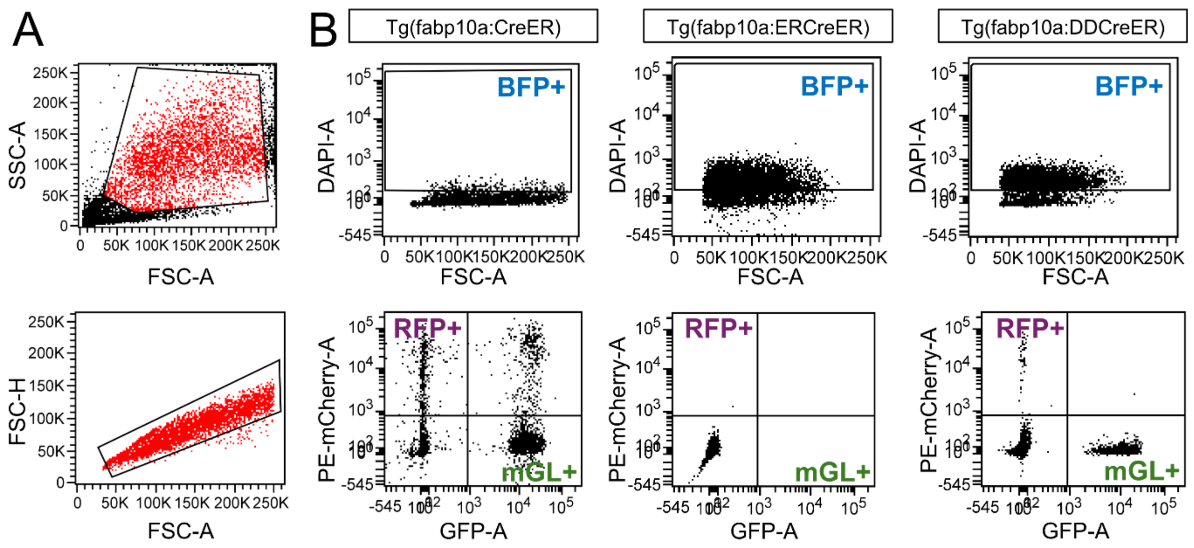


**Supplementary Figure 2: Flow cytometry analysis of background recombination in hepatocytes from different transgenic zebrafish lines.**

Representative FACS plots showing hepatocyte gating and fluorescent reporter expression in livers from 3 mpf *Tg(fabp10a:CreER)*, *Tg(fabp10a:ERCreER)*, and *Tg(fabp10a:DDCreER)* fish crossed with the multicolor reporter line *Tg(fabp10a:BB-NTR)*. No 4-hydroxytamoxifen (4-OHT) treatment was performed for any sample. **(A)** Representative forward scatter (FSC) vs. side scatter (SSC) plots used to exclude debris and FSC-A vs. FSC-H plots used to select single cells. **(B)** Top - The default mTagBFP2 signal (BFP+) is detected using excitation at 405 nm (DAPI). Bottom - mGL and mCherry are detected using excitation at 488 nm (GFP) and 561 nm (mCherry), respectively.

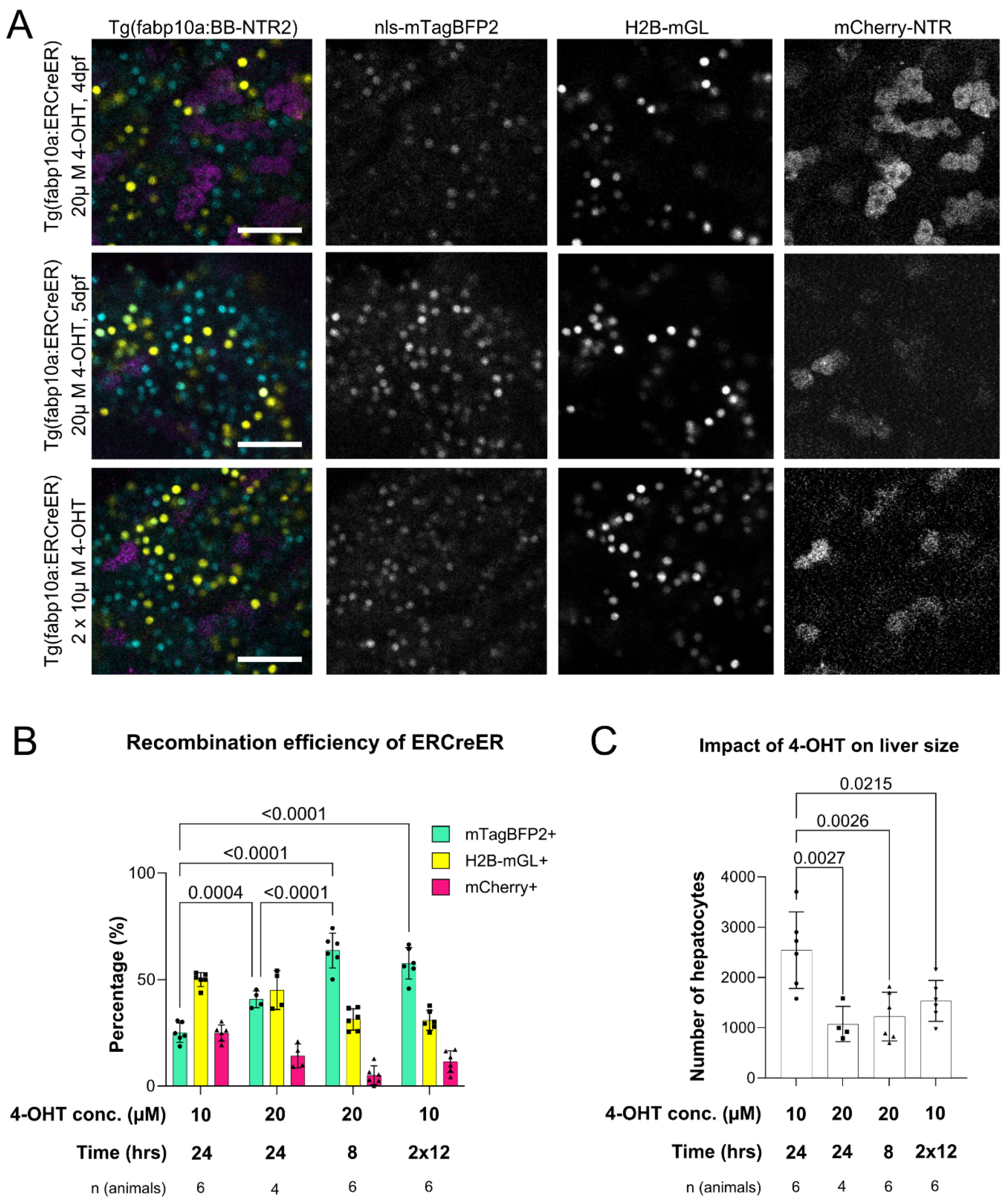


**Supplementary Figure 3: High concentrations or prolonged exposure to 4-OHT do not enhance ERCreER recombination efficiency.**

**(A)** Confocal images of livers from 10 dpf *Tg(fabp10a:BB-NTR2)*; *Tg(fabp10a:ERCreER)* zebrafish larvae following different 4-hydroxytamoxifen (4-OHT) treatments: 20 μM from 4 dpf for 24 hours, 20 μM from 5 dpf for 8 hours, or two separate 12-hour treatments with 10 μM 4-OHT, administered at 4 dpf and again at 5 dpf. Fluorescent markers indicate default (nls-mTagBFP2), and recombined (H2B-mGL and mCherry-NTR) hepatocytes. Scale bar: 50 μm. **(B)** Quantification of recombination efficiency, shown as the percentage of hepatocytes expressing each fluorescent protein (mTagBFP2, H2B-mGL, or mCherry) under the indicated 4-OHT treatment conditions. Standard treatment (10 μM 4-OHT, 4 – 5 dpf) from *Tg(fabp10a:BB-NTR)*; *Tg(fabp10a:ERCreER)* is shown as reference. Bars represent Mean ± SD. Significance was determined using Two-way ANOVA (multiple comparisons) followed by Tukey's multiple comparisons test. Number of animals analyzed per group is indicated below each bar. **(C)** Total number of hepatocytes in the left liver lobe under each treatment condition, as a proxy for liver size. Bars represent mean ± SD. Statistical comparisons were made using Two-way ANOVA followed by Tukey's multiple comparisons test. Number of animals analyzed per group is indicated below each bar.
